# A Prefusion Form of Herpes Simplex Virus 1 gB has a Distinct Antigenic Signature

**DOI:** 10.64898/2026.04.12.718011

**Authors:** Albina O. Makio, Prashanta Dutta, Jin Liu, Anthony V. Nicola

## Abstract

Herpes simplex virus (HSV) fusion and entry is mediated by a cascade of interactions among gB, gD and gH/gL. The gB homotrimer undergoes a conformational transition from a metastable prefusion state to a more stable postfusion form, driving fusion of the viral envelope and a host cell membrane. An H516P mutation in gB domain III restricts formation of the extended core alpha helix and constrains gB in a prefusion state. Several prefusion gB structures have been determined that contain this mutation. We assessed the antigenic reactivity of gB H516P by quantitative immunodotblot using a panel of gB-reactive monoclonal antibodies All antibodies tested bound to both prefusion (H516P) gB and wild type gB. Antibodies tested to gB domains II, IV and V exhibited differential binding to H516P gB compared to wild type gB. The results suggest that gB H516P has a distinct antigenic profile. The antigenic signature of H516P may be useful as a rapid indicator of prefusion forms of gB. The low pH environment of the cellular endosome is a cell-specific factor for HSV entry and triggers antigenic changes in gB. The step at which low pH impacts gB refolding to execute fusion is not well-understood. The results suggest that gB H516P undergoes wild-type-like conformational changes in gB domains I and V triggered by low pH. We propose that pH acts on an early stage of gB fusion function, prior to extension of the domain III core helix.

## Introduction

Herpes simplex virus (HSV) is a ubiquitous human virus of medical importance. There is no HSV vaccine (1). Herpes simplex virus 1 (HSV-1) fusion is coordinated and highly regulated by glycoprotein B (gB), gD, and the gH/gL heterodimer (2, 3). gB is a class III viral fusion protein conserved among herpesviruses. Herpesviral gB is a homotrimeric transmembrane protein that a contains a set of bipartite hydrophobic fusion loops (4-6). The gB ectodomain consists of several folded domains. Domain I contains the fusion loops. Domain II contains a pleckstrin homology domain. Domain III forms a central helical bundle that is extended during fusion. Domain IV forms a crown. Domain V connects Domain IV to the transmembrane domain. During fusion, gB undergoes significant domain rearrangement transitioning from a metastable prefusion state to a stable postfusion structure, accomplishing the merger of the viral and host membranes (7, 8). HSV-1 entry proceeds via a low pH endosomal pathway in a cell-specific manner (9-11). Low pH triggers changes in the antigenic conformation of gB (7, 12-14).

Characterizing prefusion structures and the ability of antibodies to bind to them is important for developing subunit vaccine strategies (15-18). Increased understanding of fusion protein conformational changes can lead to development of antiviral therapeutics such as fusion inhibitors(19, 20). The postfusion form of gB is highly stable and resistant to perturbations (21-23). Recently, several prefusion HSV gB structures were determined and analyzed by stabilizing the prefusion conformation via altering gB amino acid sequence (8, 24, 25). H156P in HSV gB domain III is a key mutation for stabilizing prefusion gB and is present in several of the determined structures. The single H516P point mutation is thought to prevent extension of the core domain III helix in prefusion gB, in turn preventing formation of an extended gB intermediate that makes fusion loop contact with the target membrane. Stabilization of prefusion gB has permitted analysis of its potential as a vaccine immunogen and as a target for antiviral drug development (8, 24-27). In this short report, we use a panel of antibodies to define the antigenic reactivity and low-pH triggered conformational change in prefusion gB H516P. Our results suggest that gB H516P has a distinct antigenic signature and has an altered trimeric conformation relative to wild type. The antigenic conformation of gBH516P was altered by low pH similar to wild type, suggesting that low pH acts at an early stage of gB conformational change.

## Results

### A single amino acid substitution in gB domain III impairs fusion in a virus-free cell-cell reporter assay

We tested the fusion activity of a pre-fusion arrested form of gB. Current models of the HSV-1 fusion mechanism are based largely on results from reporter assays of cell–cell fusion. Cells expressing HSV-1 gB, gD, gH, and gL are added to target cells bearing host cell receptors for HSV-1. Fusion of the two cell populations is indicated by a reporter, such as luciferase. In the luciferase cell-cell fusion assay, gB H516P exhibited ∼50% reduced fusion activity compared to wild type gB (**Fig. 1A**). gB H516P was similarly fusion-defective in a light microscopy-based syncytium assay (28). By CELISA, gB H516P exhibited less surface cell expression relative to wild type, but this difference was not significant (**Fig. 1B**). This suggests that the defective fusion function of gB H516P in the reporter assay was not a result of insufficient cell surface expression.

**Figure 1.**
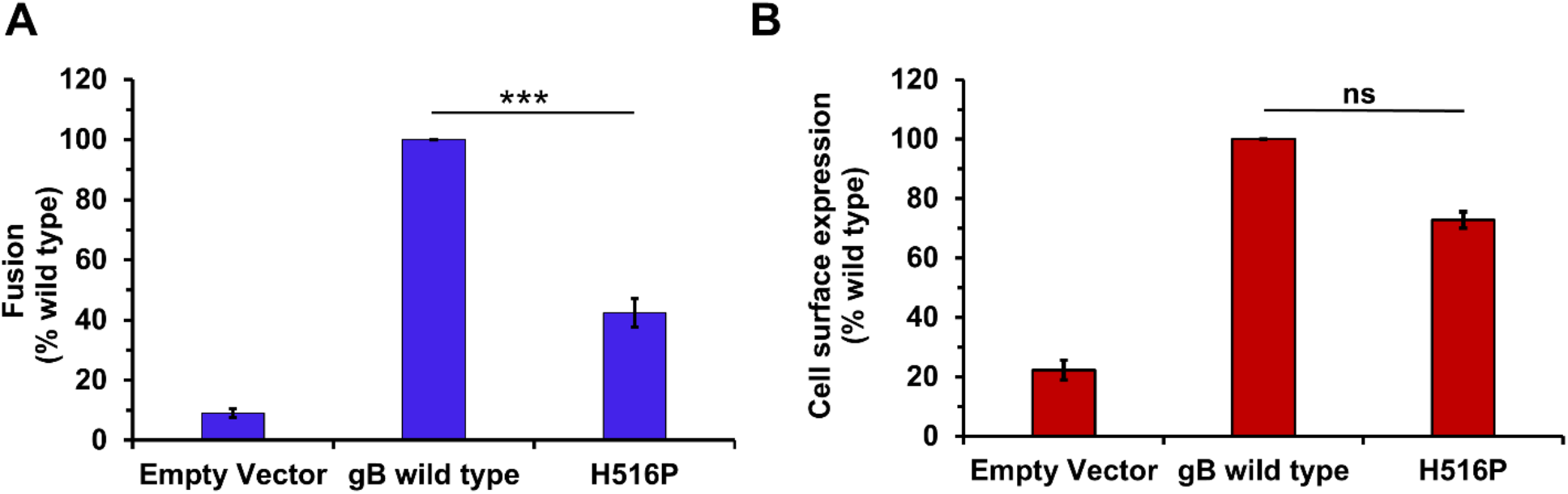
HSV-1 gB H516P is defective for fusion activity as measured by a luciferase reporter assay. **(A)** CHO-K1 effector cells transiently expressing wild type gB or gB H516P and wild type gD, gH, and gL, and T7 polymerase were co-cultured for 18 h with CHO-HVEM cells transiently expressing the luciferase gene. Luciferase-induced luminescence was quantitated as a measure of fusion. **(B)** Detection of gB on the cell surface by CELISA. CHO-K1 cells were transfected with plasmids encoding HSV-1 gB or empty vector for 24 h. Cells were fixed with paraformaldehyde and then incubated with anti-gB monoclonal antibodies H126, H1359, and H1817. HRP-conjugated Protein A was added, followed by ABTS substrate. Absorbance was read at 405 nm. Data was normalized to wild type gB set to 100% for both experiments. Results are the means of three independent experiments. *, *p* < 0.05; ns, not significant, Student’s *t-*test.

### gB H516P is antigenically distinct from wild type gB at epitopes across multiple domains

Understanding of prefusion and postfusion forms of viral fusion proteins provide start and end points for defining fusion-driving conformational changes. Engineered prefusion forms of viral glycoproteins can serve as effective vaccines (15, 18, 29). To describe the antigenic conformation of prefusion-arrested gB and distinguish it from wild type, we used a quantitative dot (**Fig. 2**). Transfected cell lysate amounts for wild type gB or H516P gB that reacted equivalently with gB-specific rabbit polyclonal antibody R68 were blotted onto nitrocellulose membrane. Blots were probed with a panel of gB monoclonal antibodies (**Fig. 2B-H**) and quantitated (**Fig. 2I**). The MAbs tested here react with both prefusion gB and postfusion gB. The differential extent to which the MAbs react to one form or the other is not known.

**Figure 2.**
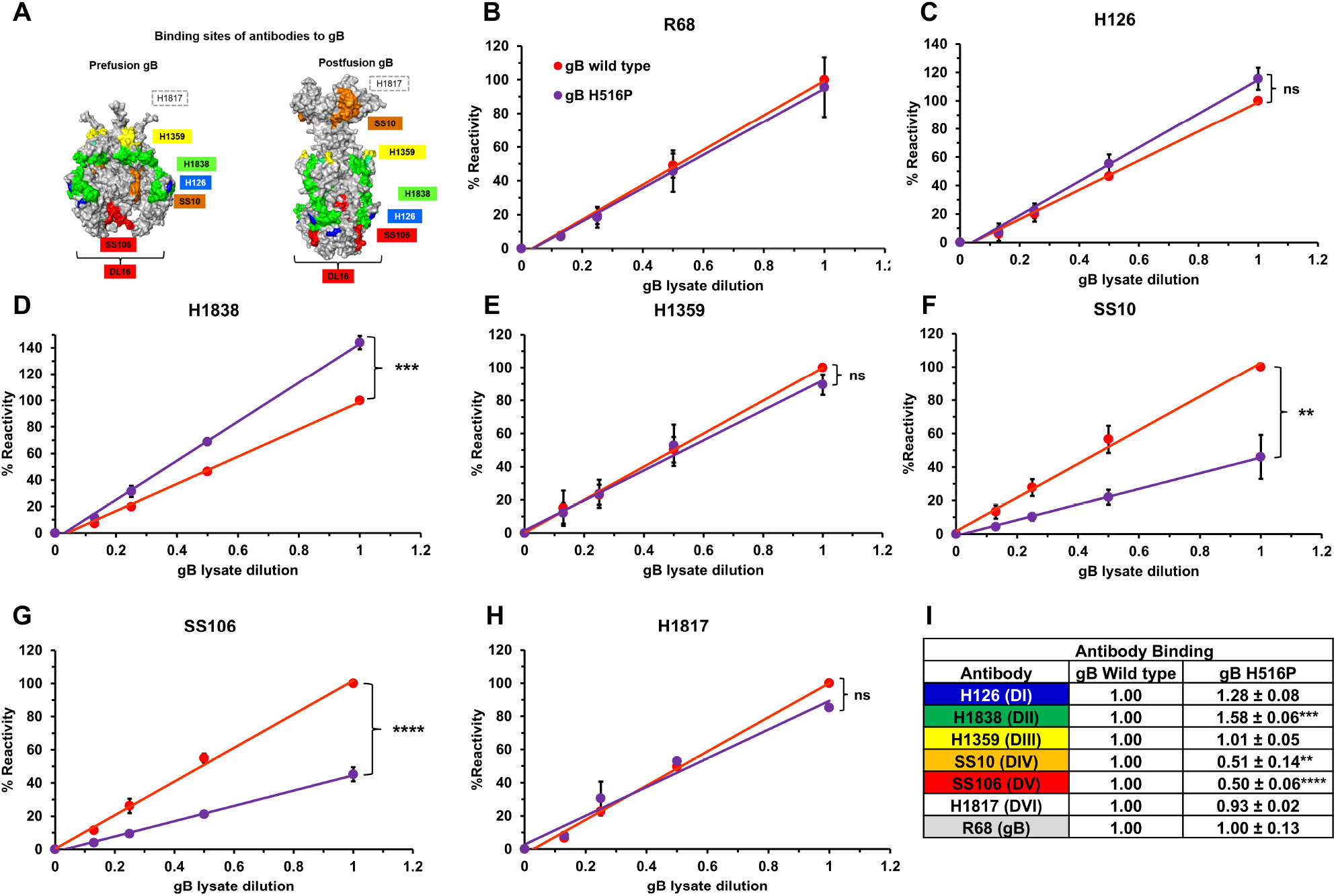
Antibody binding to a prefusion constrained gB. **(A)** Space-filling rendering (PyMOL) of prefusion gB (PDB 6Z9M) (28) and postfusion gB (PDB 2GUM) ectodomain trimer (4). Epitopes of monoclonal antibodies used in this study are highlighted in color. The MAb H126 epitope includes residue 303 in gB domain I (blue) (4, 30). The MAb H1838 epitope maps to residues 391 – 410 in domain II (green) (31). The H1359 epitope maps to residue 487-505 in domain III (yellow) (32). The SS10 epitope maps to residues 640 – 670 in domain IV (orange) (31). The SS106 epitope maps to residue 697 – 725 (33) in domain V (red). The MAb DL16 epitope is trimer-specific and resides in domain V (34). The MAb H1817 epitope maps to residues 31 – 43 in gB domain VI (white) (31), which is unresolved in the structure. **(B-H)** gB H516P is antigenically distinct from wild type gB at domains II, IV, and V. Lysates of CHO-K1 cells expressing wild type gB (red) or H516P gB (purple) were bound to nitrocellulose membrane then probed with antibodies to gB. The appropriate fluorescent-conjugated secondary antibody was added for 20 min. Images were generated with Azure Biosystems imager. Rabbit polyclonal antibody R68 was used to standard loading. Reactivity of the antibodies was quantitated relative to the wild type by densitometry using Image Studio. The values shown represent antibody reactivity relative to wild type gB set to 100%. Results are the means and standard deviations of three independent experiments. ** *p* < 0.005; *** *p* < 0.001, Student’s *t-*test. **(I)** Summary of the antigenic analysis of gB H516P. The slopes of the lines of best fit for H516P gB relative to wild type (panels B-H) were calculated and used to determine fold changes, which were normalized to 1.0.

Monoclonal antibodies H126, H1359 and H1817, which recognize epitopes in gB domains I, III and VI, respectively, reacted comparably with wild type and H516P gB **(Fig. 2C, E, H)**. This suggests similar exposure of these epitopes in prefusion and postfusion gB. In contrast, MAb H1838 to a domain II epitope exhibited increased reactivity with H516P **(Fig. 2D)**. Conversely to H1838, MAbs SS10 and SS106, to domains IV and V, respectively, had demonstrably less reactivity with H516P relative to wild type **(Fig. 2F, G)**. The results suggest that H516P has a distinct antigenic profile compared to wild type gB, exemplified by epitopes in domains II, IV and V. The antigenic signature of H516P may be useful to rapidly identify prefusion forms of gB without the need for structure determination. Results were consistent with stabilization of a distinct prefusion-like conformation of gB.

### gB H516P undergoes low pH-induced changes in antigenic conformation

The acidic intracellular milieu is an important factor in herpesvirus entry, often in a cell-specific manner. Low pH triggers changes in the antigenic conformation of HSV-1 gB (7, 12, 23, 35-37). Low pH is not sufficient to trigger conversion of gB to the postfusion conformation nor is it sufficient to trigger fusion (7, 38, 39). The precise point at which low pH acts on gB during the fusion cascade is not known. gB H516P is fusion-defective due to faulty extension of the core domain III helix, resulting in failure to form an extended intermediate gB conformation that permits fusion loop contact with the apposing membrane (8).

We measured the effect of mildly acidic pH on the conformation of gB H516P to address the stage at which pH acts on the HSV fusion process. Transfected cell lysates expressing wild type or H516P gB were exposed to pH ranging from 7.4 to 5.0 and assessed by immunodot blot at neutral pH. gB conformational change is defined as a reduction in antibody reactivity > 50%. For WT and H516P gB, the results suggested successful wild type-like conformational change in gB domains I and V. Following pH 5.0 treatment, there was a > 50% reduction in domain I MAb H126 binding to WT gB and to H516P gB (**Fig. 3A**). For MAb SS106 against gB domain V, there was a > 50% reduction in binding to both WT gB and H516P (**Fig. 3B**). Control MAb SS10 targets a domain IV epitope that is unaffected by low pH. SS10 reactivity with gB H516P remained unaltered following pH treatment (**Fig. 3C**). These results suggest that gB H516P undergoes wild-type-like conformational changes triggered by low pH. The findings further suggest that pH acts on an early stage of gB’s fusion function, prior to the extension of the central helix and contact of fusion loops with the target membrane.

**Figure 3.**
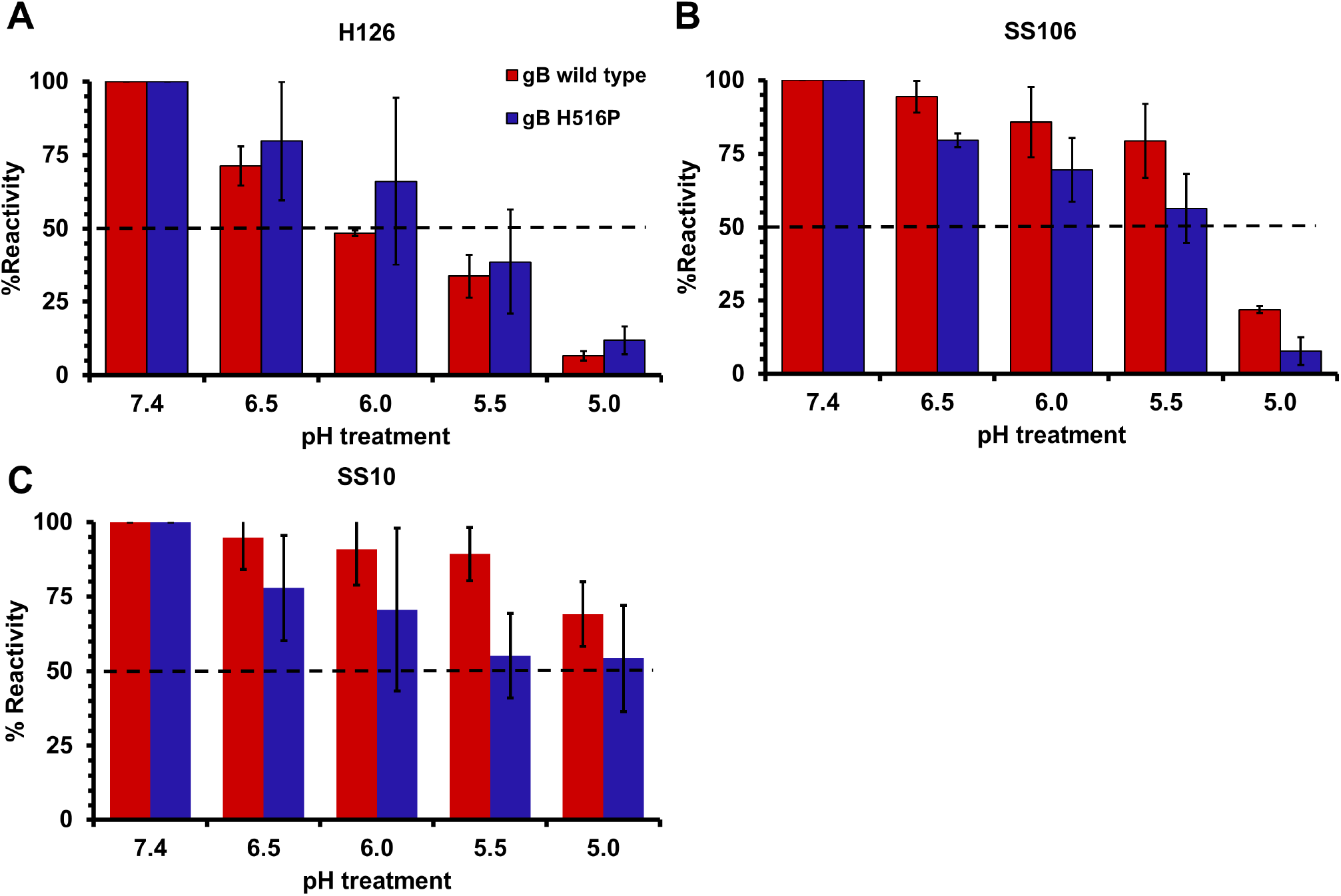
Low pH triggers conformational changes in domain I and V of H516P gB. Lysate from CHO-K1 cells expressing wild type gB or H516P gB was treated at a range of pH values (7.4 to 5.0) for 10 min at 37°C. Samples were blotted to nitrocellulose membranes and probed at neutral pH with gB MAbs targeting epitopes undergoing conformational change in **(A)** Domain 1, H126; **(B)** Domain V, SS106; and epitope not undergoing conformational change in **(C)** Domain IV, SS10 at neutral pH. Fluorescent-conjugated secondary antibody was added for 20 min. Images were generated with Azure Biosystems imager. Reactivity of three independent experiments was quantified by densitometry. The pH 7.4-treated samples were set to 100%.

### The trimeric conformation of gB H516P is distinct from wild type

The HSV-1 gB trimer is comprised of non-covalently linked monomers. We implemented two approaches to assess the oligomeric state of gB H516P. First, we compared denaturing and “native” PAGE (40) analysis of transfected cell lysates. Standard denaturing SDS-PAGE and western analysis of gB results in detection of predominantly monomeric gB, as gB trimers are disrupted by heat and SDS (41). Wild type and H516P gB monomers migrated at ∼115 kDa on denaturing SDS-PAGE, followed by western blot (**Fig. 4A**). This suggested equivalent levels of protein expression for the two gBs. For native PAGE and western blot analysis of gB, unheated samples of gB that are treated with low concentration of SDS permit detection of both trimeric and monomeric gB. Trimeric species of wild type gB were detected at >225 kDa following native PAGE and western blot, as expected. Notably, little to no gB H516P trimer was detected under the native conditions (**Fig. 4A**). Secondly, we determined reactivity of gB H516P with MAb DL16, a trimer-specific antibody to gB domain V. gB H516P exhibited diminished reactivity with MAb DL16 relative to wild type (**Fig. 4B**). The results suggest that the gB H516P has a trimerization defect or that the trimer is unstable.

**Figure 4.**
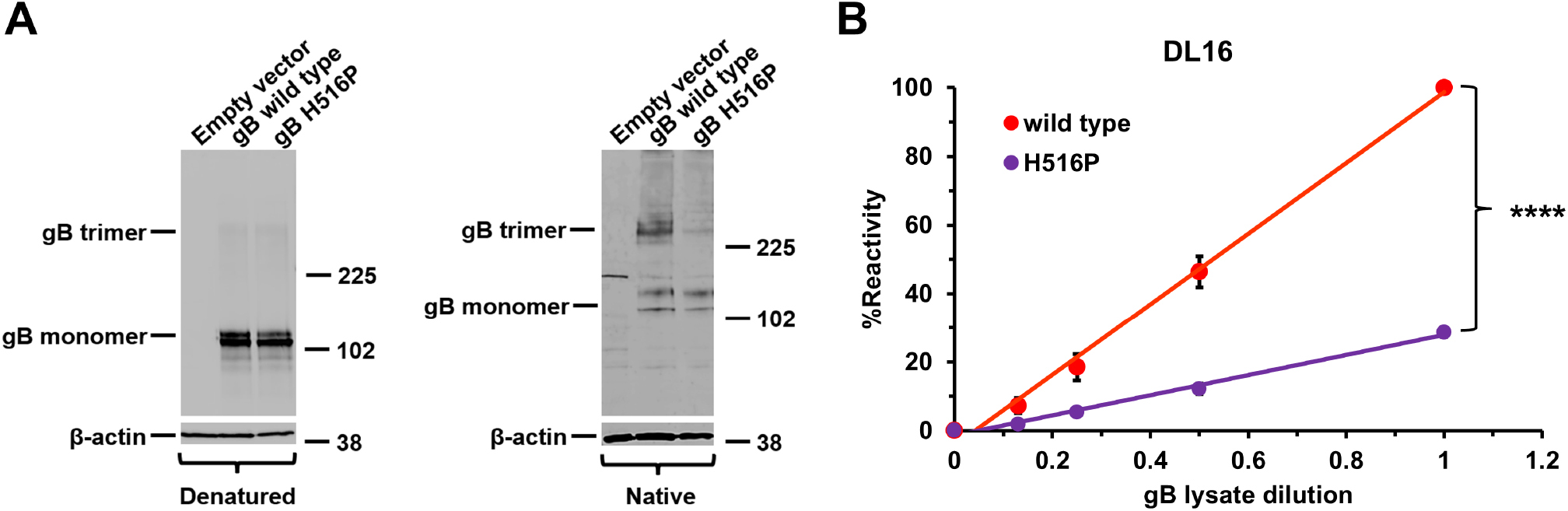
Less gB H516P trimer is detected relative to wild type gB. **(A)** CHO-K1 cells were transfected with plasmids encoding wild type gB or H516P gB. Cell lysates were either prepared in 2% SDS sample buffer with reducing agent and heated (denatured conditions), and then resolved by SDS-PAGE (left panel) or prepared with 0.2% SDS, no reducing agent, no heating (“native” conditions) and resolved by SDS-PAGE (right panel). Western blots were probed with anti-gB polyclonal antibody R68. Molecular weight standards (kDa) are indicated to the right. β-actin was included as a loading control. **(B)** Lysates of CHO-K1 cells expressing wild type gB (red) or H516P gB (purple) were bound to nitrocellulose membrane then probed with MAb DL16. Anti-mouse fluorescent-conjugated secondary antibody was added for 20 min. Images were generated with Azure Biosystems imager. Reactivity of DL16 was quantitated relative to the wild type by densitometry using Image Studio. gB wild type, 1.00; gB H516P, 0.31+/-0.01****. The values shown represent antibody reactivity relative to wild type gB set to 100%. Results are the means and standard deviations of three independent experiments. *** *p* < 0.0001, Student’s *t-*test.

## Discussion

The several prefusion gB structures that have been resolved recently represent a critical start point for elucidation of herpesviral membrane fusion mechanisms (8, 24, 25, 42-45). Here, we show that a prefusion form of HSV-1 gB containing a histidine to a proline substitution at residue 516 is impaired for fusion function as determined by a virus-free cell-cell fusion assay **(Fig. 1)**. The antigenic structure of H516P gB exhibits a unique signature distinct from wild type at domains II, IV, and V **(Fig. 2)**. Similar to wild type gB, H516P gB undergoes pH-induced conformational changes when subjected to mildly acidic pH **(Fig. 3)**, suggesting that pH acts on gB early, before the formation of the extended alpha helix in domain III. Lastly, the H516P gB trimer is detected poorly compared to wild type despite similar protein expression levels **(Fig. 4)**.

Wild type gB when expressed alone favors the postfusion conformation (4, 8, 21). Each of the MAbs tested here react with both prefusion gB (H516P) and postfusion gB (wild type). This is in line with the recent finding that a significant antigenic surface area remains unchanged in prefusion and postfusion gB (24). Binding of H1838, SS10, SS106, and DL16 antibodies to prefusion gB was measurably distinct from wild type **(Fig. 2)**. Thus, antigenic analysis is a simple and efficient means to distinguish prefusion gB from postfusion gB.

Prefusion-arrested subunit vaccines have proven effective for several viruses (15-18, 27, 29). Stabilization of prefusion gB has allowed assessment of its immune characteristics. A llama nanobody was isolated that is specific to prefusion gB and neutralizes HSV-1 in cell culture (25). An antibody generated against the prefusion gB-stabilized ectodomain was specific for prefusion gB and neutralized HSV-1. However, sera from mice immunized with prefusion stabilized gB ectodomain did not neutralize HSV-1 (24). In the present study, H1838 was the only antibody tested that bound significantly better to prefusion gB **(Fig. 2D, I)**. H1838 is a virus-neutralizing antibody (32). It is tempting to speculate that the mechanism of H1838 neutralization is via blocking the transition of prefusion gB to postfusion gB. An mRNA vaccine study in mice suggested that gB H516P exhibited immunogenicity similar to wild type (26). Neutralizing monoclonal antibody 16F binds to a broad-spectrum herpesvirus epitope in gB domain I and is thought to bind both prefusion and postfusion gBs (27).

Under native PAGE conditions, the H516P gB trimer is virtually undetectable. Using an independent approach, a trimer-specific antibody exhibited reduced reactivity with H516P **(Fig. 4)**, also suggesting the trimer is less detectable. Impaired gB trimerization or reduced trimer stability might contribute to H516P gB’s impaired fusion function. Mildly acidic pH induces antigenic changes in domains I and V of wild type gB that are thought to be important for fusion (7, 12, 23, 37, 46). H516P gB undergoes pH-induced conformational changes in domains I and V similar to wild type **(Fig. 3)**. gB fusion loop mutants are also defective in fusion and undergo wild type-like antigenic changes in response to low pH (5, 23). Because H516P gB is functionally impaired for fusion yet still responds to acidic pH, these results suggest that low pH acts on an early stage of gB’s fusion function, prior to formation of the proposed extended intermediate that requires hinge flexibility in domain III.

## Materials and Methods

### Cells

Chinese hamster ovary (CHO-K1) cells (American Type Culture Collection (ATCC), Manassas, VA, USA) were propagated in Ham’s F12 nutrient mixture (Gibco/Life Technologies, Grand Island, NY, USA) supplemented with 10% fetal bovine serum (FBS) (Atlanta Biologicals, Atlanta, GA, USA). CHO-K1 cells expressing the human herpesvirus entry mediator (CHO-HVEM or M1A cells) and containing the *lacZ* gene under control of an HSV-inducible promoter (47) were obtained from G. Cohen and R. Eisenberg, University of Pennsylvania. CHO-HVEM cells were propagated in Ham’s F12 nutrient mixture supplemented with 10% FBS, 150 µg/mL puromycin (Sigma Aldrich, St. Louis, MO, USA), and 250 µg/mL G418 sulfate (Sigma Aldrich, St. Louis, MO, USA). Cells were subcultured in non-selective medium for at least two passages before use in experiments.

### Antibodies

Anti-HSV-1 gB mouse monoclonal antibodies H126 (4, 30), H1359 (32), and H1817 (32, 48) were purchased from Virusys, Taneytown, MD, USA. MAb H1838 mouse ascites fluid (32) was provided by L. Pereira, University of California, San Francisco. Anti-gB mouse monoclonal antibodies DL16 (oligomer-specific), SS10, and SS106, (48), and anti-gB rabbit polyclonal serum R68 (49) were provided by G. Cohen and R. Eisenberg, University of Pennsylvania. Anti-beta-actin monoclonal antibody AC-74 was purchased from Sigma (St. Louis, MO, USA).

### Construction of HSV-1 gB with a point mutation (H516P) in domain III

The construction of pAM24, which contains HSV-1 KOS wild type gB sequence (50) with *NsiI* and *EcoRI* flanking sites upstream and downstream of gB, respectively, in a pCAGGS/MCS backbone (51, 52) was previously described (53). A single nucleotide point mutation designed to change histidine to proline at residue 516 was introduced in the gB sequence using the Q5 Site-Directed Mutagenesis Kit (New England Biolabs, Ipswich, MA, USA). The mutagenic primer pair consisted of a sense primer 5’ ATC CAG GGG CCC GTG AAC GAC – 3’ and an antisense primer 5’ GTG GTT GTA GGT AAA CTG CAG – 3’. The generated plasmid construct was digested with *NsiI, Ase I*, and *EcoRI* enzymes, and the pCAGGS/MCS backbone was digested with *NsiI* and *EcoRI* enzymes. The digestion products were gel-purified with a QiaQuick Gel Extraction Kit (Qiagen). The gB gene fragment and pCAGGS/MCS DNAs were then ligated using Instant Sticky-End Ligase (New England Biolabs) and transformed into competent *E. coli* DH5-α cells (New England Biolabs), generating pAM28 (gB H516P). All restriction enzymes were from New England Biolabs. All plasmids were sequence-verified.

### Luciferase reporter assay for cell-cell fusion

CHO-K1 (effector) cells were transfected with plasmids encoding T7 RNA polymerase (pCAGT7) (54), HSV-1 KOS wild type gB (pAM24) or gB H516P (pAM28), HSV-1 KOS gD (pPEP99), gH (pPEP100), and gL (pPEP101). Plasmid series pPEP provided by P. Spear, Northwestern University (55). CHO-HVEM (target) cells were transfected with a plasmid encoding the firefly luciferase gene under the control of the T7 promoter (pT7EMCLuc) (55). All cells were transfected for 6 h at 37°C in OptiMEM (Life Technologies, Grand Island, NY, USA) using the Lipofectamine 3000 system (Invitrogen, Carlsbad, CA, USA). A single cell suspension of target cells was added to effector cell monolayers and co-cultured in Ham’s F12 medium for 18 h at 37°C. Cells were lysed using the Promega Luciferase Assay System (Madison, WI, USA), and lysates were frozen and thawed. Substrate was added to cell lysates and immediately assayed for light output (luciferase activity; fusion) using a BioTek Synergy Neo microplate luminometer (Santa Clara, CA, USA).

### Analysis of cell surface-expressed gB by CELISA

CHO-K1 cells in 96-well plates were transfected with plasmids encoding HSV-1 KOS wildtype gB (pAM24) or gB H516P (pAM28), or empty vector (pCAGGS/MCS) with Lipofectamine 3000 (Invitrogen, Carlsbad, CA, USA). Cells were cultured for 18 h and then fixed in 4% paraformaldehyde. Fixed cells were blocked with 3% BSA in PBS for 2 h. Anti-gB monoclonal antibodies H126, H1359, or H1817 in 3% BSA in PBS were added overnight at 4°C. Protein A conjugated to horseradish peroxidase (Invitrogen) was added for 2 h at room temperature. Substrate 2,2′-Azinobis [3-ethylbenzothiazoline-6-sulfonic acid]-diammonium salt (ABTS; Thermo Fisher Scientific) was added, and absorbance was measured at 405 nm using a BioTek microplate reader.

### Antigenic analysis of gB

CHO-K1 cells transfected with either pAM24 (wild type gB) or pAM28 (gB H516P) were rinsed with PBS, then lysed in either 2% 3-[(3-cholamidopropyl)dimethylammonio]-1-propanesulfonate (CHAPS) (Sigma Aldrich, St. Louis, MO, USA) detergent in HEPES-buffered saline (HBS) (ACROS Organics, New Jersey, USA) with Halt protease inhibitor (Rockford, IL, USA). Detergent lysates were pelleted at 20,000 x *g* for 10 min at 4°C. Supernatant was collected and stored at -80°C. Supernatants (∼ 1.3 × 10^5^ cell equivalents) were diluted two-fold in serum-free, bicarbonate-free DMEM with 0.2% bovine serum albumin (BSA) and 5 mM each: HEPES, MES (morpholineethanesulfonic acid), and sodium succinate and blotted directly to a nitrocellulose membrane using a Minifold dot blot system (Whatman) (35). Membranes were blocked with 5% milk in PBS containing 0.2% Tween 20 (blocking buffer) and then incubated with antibodies to gB in blocking buffer at 4°C overnight. After incubation with goat anti-mouse or goat anti-rabbit Alexa Fluor 647 secondary antibodies (Thermo Fisher Scientific, Rockford, IL, USA) for 20 min in blocking buffer, images were obtained with an Azure Biosystems imager C400 (Dublin, CA, USA). Densitometry was performed with Image Studio (Version 6.1) (LI-COR Biosystems, Lincoln, Nebraska, USA). Means of three independent experiments with standard deviation were calculated. Wild type gB reactivity with each antibody was normalized to 1.00.

### Low pH-induced changes in antigenic conformation of gB

CHO-K1 cells transfected with either pAM24 (wild type gB) or pAM28 (gB H516P) were rinsed with PBS, dislodged by scraping into HBS with Halt protease inhibitor. Cells were pelleted at 1,250 x *g* for 5 min at 4°C. Supernatant was discarded, and cells were resuspended in HBS with Halt protease inhibitor. Cells were subjected to four cycles of sonication. Each cycle was 1 min with 10 sec pulses (50 decibels) with a Sonic Dismembrator model 705 (Thermo Fisher Scientific, Waltham, MA, USA). Samples were rested on ice between cycles. Sonicated lysates were pelleted at 20,000 x *g* for 5 min at 4°C. Supernatant was collected and stored at -80°C. gB-expressing lysates (approximately 1.3 × 10^5^ cell equivalents) were diluted in serum-free, bicarbonate-free DMEM with 0.2% bovine serum albumin (BSA) and 5 mM each of HEPES, MES (morpholineethanesulfonic acid), and sodium succinate and adjusted with a pretitrated amount of 1.2 N HCl for 10 min at 37°C to achieve final pHs ranging from 7.4 to 5.0. Samples were blotted directly to a nitrocellulose membrane using a Minifold dot blot system. Membranes were blocked with blocking buffer.

Membranes were incubated with anti-gB monoclonal antibodies in blocking buffer at neutral pH. Followed by incubation with goat anti-mouse Alexa Fluor 647 antibody in blocking buffer. The membrane was imaged with an Azure Biosystems c400 imager and quantified by densitometry with Image Studio.

### SDS – PAGE and western blot analysis of gB

Transfected cell lysates were either treated with modified Laemmli sample buffer with 0.2% sodium dodecyl sulphate (SDS) (low SDS) and no reducing agent (“native” conditions) (56) without heating or added to Laemmli sample buffer containing 2% SDS, 200 mM dithiothreitol (DTT) and heated to 85°C for 5 min (denaturing conditions). Proteins were separated on 4–20% Tris-glycine gels (Invitrogen, Carlsbad, CA, USA). Gels were transferred to a nitrocellulose membrane and then blocked with 5% milk in PBS containing 0.2% Tween 20 (blocking buffer) for 20 min. Membranes were probed overnight with primary polyclonal antibody to HSV-1 gB (R68) (49) in blocking buffer. Fluorescent-conjugated anti-rabbit secondary antibody was added for 20 min in blocking buffer.

Images were obtained using an Azure Biosystems imager and analyzed by densitometry (Image Studio).

## Declarations

### Ethics approval and consent to participate

Not applicable.

### Consent for publication

Not applicable.

### Availability of data and materials

All relevant data are contained within the paper. Reagents and data are available upon reasonable request to the corresponding author.

### Authors’ contributions

Conceptualization, A.O.M., P.D., J.L. and A.V.N.; methodology, A.O.M and A.V.N.; investigation, A.O.M.; formal analysis, A.O.M.; writing original draft preparation, A.O.M; writing—review and editing, P.D., J.L. and A.V.N.; supervision, A.V.N.; funding, P.D., J.L. and A.V.N. All authors have read and agreed to the published version of the manuscript.

## Acknowledgements

We thank Gary Cohen, Roselyn Eisenberg, Lenore Pereira, and Patricia Spear for gifts of reagents.

